# Emerging combinations of climatic parameters for dengue proliferation in urban landscapes

**DOI:** 10.64898/2026.05.19.726173

**Authors:** Aditya Vaishya, Vaibhavi Patel, Yash Dahima, Lalit Shankar Chowdhury, Kaushik Jana, Bhargav Adhvaryu, Darshini Mahadevia, Chirag Shah, Subhash Rajpurohit

## Abstract

Ectotherm insects’ growth and development are dictated by temperature and humidity. Conducive habitats and the availability of resources set ideal conditions for insect population growth. Mosquitoes require water, favorable temperature, and blood meal to survive. In this research, we picked a rapidly growing megacity, Ahmedabad, in western India, to explore and establish potential linkages between disease spread and meteorological conditions. Ahmedabad, with a population of over 8 million, is experiencing changes in rain and humidity patterns, pushing the city towards changing vector-borne disease dynamics. We examined dengue cases over ten years, 2012-22, and explored their connections with two prominent climatic variables, temperature and relative humidity. Our findings indicate that stable temperature (25-27.5 °C) and humidity (> 60%) interaction is a ruling factor in spikes in dengue cases in the city. While stable temperature ranges triggers the dengue cases, RH drives the explosive phases and sustainability of such episodes. Statistically significant increasing trends in temperatures, narrowing down of the day-night temperature ranges, and increasing night temperatures provide more stable temperature regimes in a warming world thereby likely to extend the dengue season beyond the usual monsoon season.

**Plain Language Summary:** Dengue incidences have been found to be associated with mosquito population outbreaks. Every year, thousands of lives are lost due to this deadly virus spread by mosquitoes. Particularly in the Indian subcontinent, a large proportion of these cases is associated with the monsoon season and rain patterns. In recent years, there have been abrupt spikes in dengue cases across Indian cities, particularly in western India. To understand this complex interaction of viral proliferation and local environmental conditions, the last ten years of dengue case patterns have been scanned in parallel to the climate data. Our findings suggest that stable temperature windows and humidity levels above certain thresholds trigger a rise in dengue cases. While stable temperature ranges trigger dengue cases, humidity drives such episodes’ explosive phases and sustainability. Our work pinpoints specific temperature-humidity combinations and suggests that local municipal corporations use them as warning indicators to initiate preventive measures.

## Introduction

Local variations in vector-borne disease risk is influenced by climatic variables like rainfall, temperature, and relative humidity (RH) [1, 2]. Tropical climatic conditions of the Indian subcontinent suit many insect species identified as vectors for transmitting pathogens of diseases like malaria, dengue, Japanese encephalitis, kala-azar, lymphatic filariasis, and chikungunya. Approximately half of the human population is at risk of infection of a vector-borne disease [3]. The largest contributors of human vector-borne disease are transmitted by arthropod vectors, such as ticks and mosquitoes [4]. One of these diseases, which is endemic in more than 100 countries, is dengue, particularly in Asia, Africa, and the Americas. Dengue virus infects almost 390 million people worldwide yearly in tropical and subtropical regions [5]. Of total number of dengue cases worldwide almost 70% occur in Asia [5, 6]. According to recent studies, over the last two decades (2000-2019), dengue cases have increased ten-fold [7, 8]. There are new cases emerging in Europe, with Croatia and France reporting their first cases in 2010 [9-11]. Temperature is a factor that plays a vital role in the transmission of this disease [5, 6]. The favorable temperature ranges for dengue (DENV) virus transmission is 13-35°C [12-14]. Changes in the climate can influence the local environment on both micro and macro scales [15]. Frequent and sustained higher temperatures are expanding the breeding grounds and bringing along Zika, dengue viruses, and malaria parasites to these areas for the first time [16]. Intergovernmental Panel on Climate Change’s (IPCC’s) Assessment Report 6, Working Group 2 (WGII) report [17] has confirmed that the incidence of vector-borne diseases has increased and is projected to increase under all levels of (projected) warming scenarios without additional adaptation [17].

Dengue virus (DENV) is a single positive-stranded RNA virus belonging to the family Flaviviridae. Four serotypes are present, DENV-I, DENV-II, DENV-III, and DENV-IV, which are loosely antigenically distinct [18, 19]. The incidence of dengue has grown significantly in recent decades [20]. This virus is transmitted mostly by female mosquitoes of the species *Aedes aegypti* and to a smaller extent, *Aedes albopictus* [19]. *Aedes aegypti* is a domesticated species that can be found near or in houses and buildings. It lays eggs in stagnant fresh water, tap water, or rainwater stored in containers [21, 22]. These mosquitoes are also vectors of chikungunya, yellow fever, and Zika virus. DENV is frequently transported from one place to another by infected travelers. This was seen in northern Australia, which had no incidence of dengue for more than 25 years. However, outbreaks were reported after the opening of an international airport in Cairns [22].

Having an incubation period of 5 to 7 days, DENV causes severe headache, retro-orbital pain, muscle, joint and bone pain, and maculopapular rash unless asymptomatic [23]. Interestingly, the vast majority of DENV cases are asymptomatic, and when vectors are present in these new areas, there is potential for transmission [24]. There is no specific treatment for dengue or severe dengue except for managing the vital body parameters of the patients. We think that the lack of early warning mechanisms, in this case, is due to knowledge gaps in DENV biology and its interaction with climatic variables (i.e. organism and environment interactions).

In India, DENV transmission occurs in almost all the parts of the country due to climate suitability for dengue transmission except the upper part of Jammu and Kashmir, Uttarakhand, Sikkim and Arunachal Pradesh [25, 26]. Entire states of Tamil Nadu, and parts of Karnataka, Orissa, Andhra Pradesh, West Bengal, and North East and Bihar are open to dengue transmission for 10-12 months a year; parts of Gujarat (i.e. Kutch, Saurashtra coastal areas, and parts of South Gujarat) for 7-8 months and the rest of Gujarat for 4-6 months [25]. This gets even more severe during mid to late monsoon [26]. The trend of rapid increase in dengue cases during monsoon months in the last decade has also led to increased hospitalization and many death counts [26]. Over the last 20 years, number of dengue cases and the associated death toll has increased significantly across India [27] and an increase of over 70% was reported during the period 1990 - 2019 [28]. In recent years, the National Center for Disease Control (NCDC) has identified dengue outbreaks in several parts of the country (https://ncdc.mohfw.gov.in/). An unparalleled increase in dengue cases in the next few years could lead to a severe burden on the public health system and economy.

India is a subtropical country which has clear seasons. The seasonal variations get prominent as one moves from southern to northern regions of the country i.e. coefficient of variation of temperature. In Indian subcontinent, monsoon (rainy season) window is from early June to September. During this season, most of the habitats are water rich and humid with moderate to high-temperatures which fluctuates throughout the rainy season with intermittent stable periods. The monsoon season in India is crucial for the crop cycling and farmers’ lives whereas excessive rains during this period triggers disease. While it relieves the hot summers and is the essence of India’s rapidly expanding economy, it also triggers a series of water-borne and vector-borne diseases [29, 30]. The monsoon rains provide the much-needed water to the farms and recharge the aquifers and reservoirs [31, 32]. While this is good for the economy and sustainability of biodiversity, the flip side is that it triggers public health issues. The warm conditions, as well as the water and the high population density in some cities of India, provide ideal conditions for the onset of vector-borne diseases, particularly dengue. In India, it has been observed that dengue cases peak somewhere around the monsoon - post-monsoon period. Figure 1 shows the map of India with districts, colored in red, prone to dengue and having a statistically significant increasing summer monsoon rainfall trend of 20% or more over 40 years (1982-2022) continuous time-series. The data for the dengue prone districts were fetched from the National Centre for Vector Borne Diseases Control [33] and for rainfall from Prabhu and Chitale, 2024 [27].

**Figure 1:**
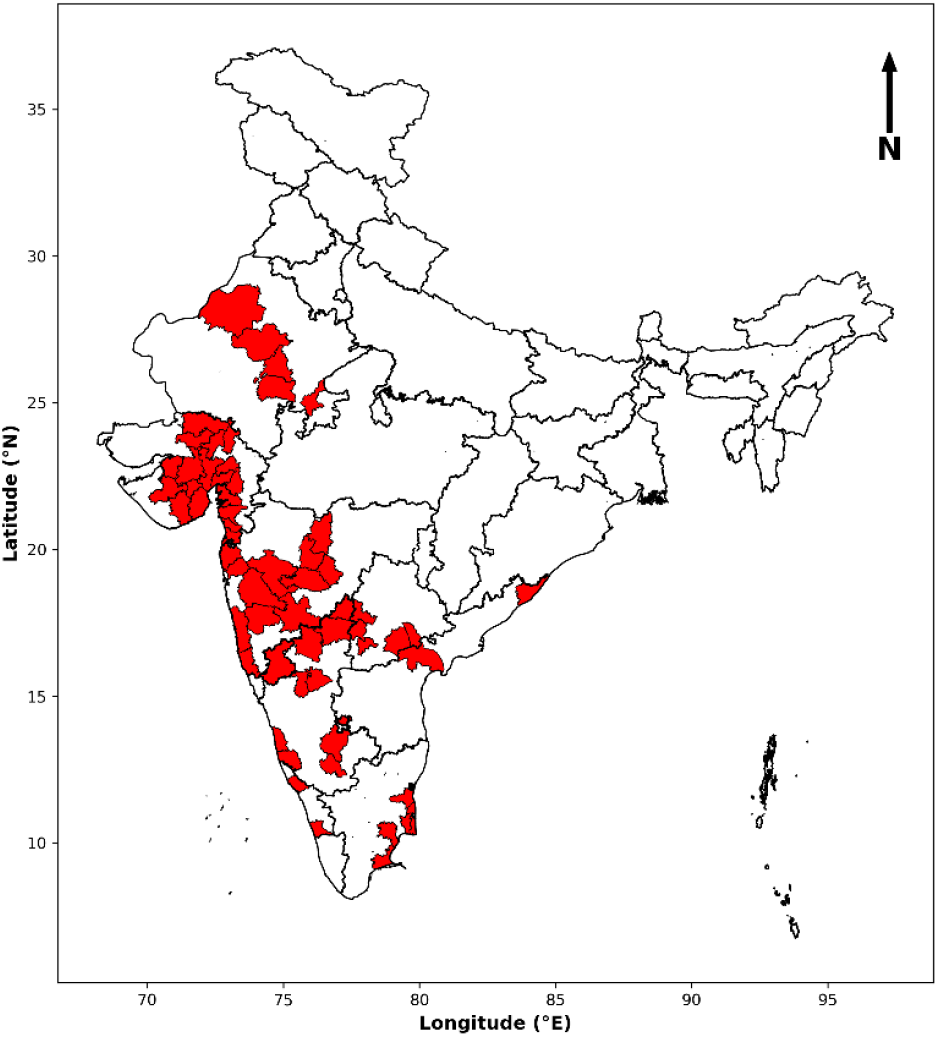
Map of India showing districts, colored in red, prone to dengue and having a statistically significant summer monsoon rainfall trend of 20% or more over 40 years (1982-2022) continuous time-series. The data for the dengue prone districts were fetched from the National Centre for Vector Borne Diseases Control [33] and for rainfall from Prabhu and Chitale, 2024 [34].

Gujarat, a state with a population of roughly 60 million people, has one of the highest cases of dengue. It is evident from Figure 1 that dengue cases in the state will likely rise in future as the summer monsoon rainfall increases. According to the National Center for Vector Borne Diseases Control, in 2019, Gujarat had over 18,000 cases, the highest of any other state in India [33]. It has a tropical monsoon climate, and except for the rainy season, it is usually hot and dry, with an average annual rainfall of over 700 mm in monsoon [35]. It is reported that the correlation coefficient between rainfall and dengue cases in the city of Ahmedabad in Gujarat showed a significant correlation at the lag of 2 months [25]. But, no studies are available that establish a link between temperatures and relative humidity with dengue cases individually or together. Within Gujarat, the city of Ahmedabad is going to be next metropolitan of the country with several urban life challenges including proper drainage, changing monsoon patterns, and dense population establishments [36, 37]. Having a population of approximately 7.6 million and a tropical monsoon climate, it has the population as well as the environmental conditions to be an ideal city to study the correlation between the climatic changes with the number of dengue cases.

Insects inhabit a wide range of habitats across the planet. Climate parameters like temperature and humidity impact insect development, growth, mortality, and behaviour [38-40]. At higher temperatures, insects grow faster, i.e., they have faster development time [41]. Also, insects face other physical constraints. For example, insects are smaller in size, exposing them to drought (i.e., larger surface area to volume ratio). Under drier conditions, small insects can easily lose body water as they balance body water through the hard cuticle [42, 43]. Collectively, temperature and humidity play a major role in insect’s fitness [44-46].

Effects of temperature and humidity on insect reproductive physiology have been explored at length but in isolation and primarily focusing on temperature only [47-51]. A few studies have also examined temperature effects on vector’s behaviour, development rate, and mortality [38-40]. Very few studies have attempted to understand the impact of *temperature-humidity* interactions on reproductive fitness [52]. On these lines, our understanding of insect vectors is even more limited.

Field-based evidence or studies are rare. Insect vectors (e.g., mosquitos) and human interactions are complicated. Females of many mosquito species blood meal on humans and other mammals to produce eggs. It’s an essential part of their reproductive physiology [53]. Understanding their role in human health and disease is complex, and a deeper understanding of the links between diseases and climate is needed [54]. Also, the impact of changing climatic conditions on dengue spread is unclear. In their recent work, Prasad et al. [55] demonstrated that the mosquito eggs rewire their polyamine and lipid metabolism to survive extended drought periods [55]. This is crucial as rapid lipid breakdown is required in larval hatching and survival upon rehydration [55]. Post-monsoon dengue cases outcomes could be possibly due to the adaptations of mosquito eggs to drier environments, however this cannot be verified with the existing data.

Here, we propose to fill this gap in our understanding of this disease by studying the interaction between dengue cases and climatic conditions in the megacity of Ahmedabad. By looking at the variation in climatic conditions, we hypothesize to see a pattern in the dengue cases with systematic changes in temperature and relative humidity. Specifically, we want to see if there exist any specific temperature and relative humidity ranges in which dengue proliferation happen in this region. Thus, we aim to establish linkages between selected climate variables and abundance in dengue cases.

## Material and methods

### Dengue cases data

The data of dengue cases in Ahmedabad, collected by the Ahmedabad Municipal Corporation (AMC), was made available for this research by the Medical Officer of Health (MoH), AMC. The collected data spans from 2012 - 2022, consisting of cases registered in government and private hospitals in Ahmedabad. The dengue cases are organized monthly for each year. Data before 2012 were not available. Besides, the number of cases relates to the ones reported and hence registered. There could be more cases in the city than registered. This study is limited to the reported and registered cases in the period stated. Further, the data obtained from the AMC related to the cases reported in the hospitals located within the boundary of the AMC. The cases are not collected at the source of origin. Further, the data pertains to the reported cases of dengue, that is, when the disease is detected, and there might be other asymptomatic carriers of the virus who remain undetected. For such an analysis sero surveys have to be undertaken in host populations, which we have not done. Hence, the study should be seen as identifier of temperature and humidity markers that warn about potential outbreak of dengue, for which local level actions need to be taken.

### Climate parameters

The meteorological parameters 2 meter air temperature and 2 meter air dew-point temperature are ERA5-Land reanalysis products provided by the European Centre for Medium Range Weather Forecast (ECMWF). The 2m air RH was derived using temperature and dew-point temperature using the formula below [56, 57]:

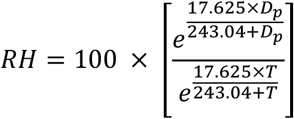

Here, *RH* is relative humidity (%), *T* is 2m air temperature (°C), and *D*_*p*_ is 2m air dew point temperature (°C). This dataset’s spatial and temporal resolution is 10 km and 1 hour, respectively. The land element of the ERA5 climate reanalysis is re-run to develop the ERA5-Land reanalysis product [58]. The global measurements and the data from models were integrated, and validated to create the reanalysis products.

The Land Surface Temperature (LST) is the satellite data product provided by the MODIS instrument onboard Terra satellite. The spatial and temporal resolution of this dataset is 1 km and 1 day, respectively. The products MOD21A1D and MOD21A1N were used for daytime and nighttime LST. A physics-based approach is applied in the MOD21 TES (Temperature/Emissivity Separation) algorithm [59] to retrieve spectral emissivity and LST at the same time from the MODIS thermal infrared bands 29, 31, and 32. To ensure accurate results, particularly at very high temperature and RH values, the TES algorithm is coupled with an enhanced Water Vapor Scaling (WVS) atmospheric correction method [60].

The final dataset consists of values of hourly 2m air temperature, hourly 2m air RH, average daytime LST, and average night time LST for the years 2012 – 2022 averaged over Ahmedabad Municipal Corporation boundary (bounded by latitude 22.9°N – 23.2°N, and longitude 72.4°E – 72.8°E).

### Mann-Kendall test

The Mann-Kendall test (MK) [61, 62] is a non-parametric statistical method and is widely used in environmental and climate studies to detect trends in time series data. The test assesses whether there is a monotonic upward or downward trend in the data which may or may not be normally distributed. The trend is said to be statistically significant if the associated p value is < 0.05. A positive value of the trend component with p < 0.05 indicates a statistically increasing trend, while a negative value of the trend component with p < 0.05 indicates a statistically decreasing trend.

## Results

Examining the meteorological data spanning from 2012 to 2022 for Ahmedabad, seasonal trends emerge in the temperature and RH patterns. The winter months experienced the lowest temperatures, with a seasonal average minimum ranging from 14.2°C to 16.1°C and a maximum ranging from 27.2°C to 29.5°C over the years, and January mostly being the coolest month. The summer months observed the highest temperatures, where the seasonal average maximum temperature was peaking between 38.0°C and 40.2°C, and May was mostly the hottest month.

Similarly, the RH levels followed seasonal patterns. Pre-monsoon months (Mar-May) tended to be drier, with RH levels hovering between a minimum of (20.0 ± 1.9) % and maximum of (63.3 ± 2.8) %, while monsoon experienced higher humidity, with RH level ranging from minimum (64.2 ± 3.3) % to maximum (90.7 ± 1.7) %. These seasonal statistics illustrate the cyclic nature of temperature and humidity variations, representing the distinctive climatic attributes of Ahmedabad across different seasons.

To identify trend in meteorological parameters, the Mann-Kendall test was performed. The native frequency of temperature and RH data was hourly. First of all, the following parameters were derived using the data: Min T, Max T, Avg T, ΔT, Min RH, Max RH, Avg RH, and ΔRH. Here, Min, Max, and Avg indicates the daily minimum, daily maximum, and daily average values of the corresponding meteorological parameters; whereas Δ shows the difference between daily maximum and daily minimum values of the corresponding variable.

Seasonal, residual, and trend components were separated for the above mentioned parameters and the Mann-Kendall test was performed on the trend components of each parameter. Table 1 shows the results of this test. All values in the table are statistically significant (p < 0.05) except Max RH. It can be observed from the table that minimum values of these meteorological parameters are increasing more rapidly than the maximum values leading to more stable temperature and RH regimes.

**Table 1:**
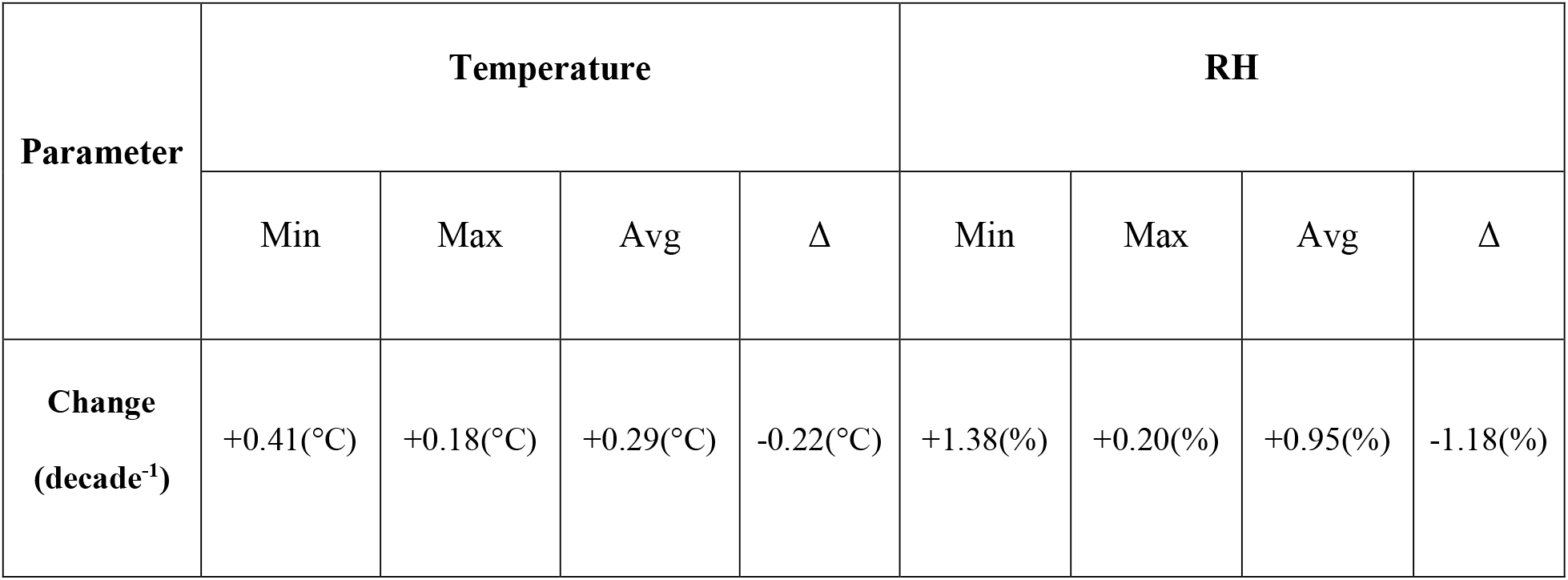
Change in the trend components of temperature and RH parameters over a decade from 2012 – 2022.

| Parameter | Temperature |  |  |  | RH |  |  |  |
| --- | --- | --- | --- | --- | --- | --- | --- | --- |
| | Min | Max | Avg | $\Delta$ | Min | Max | Avg | $\Delta$ |
| **Change**<br>**(decade<sup>-1</sup>)** | +0.41(°C) | +0.18(°C) | +0.29(°C) | -0.22(°C) | +1.38(%) | +0.20(%) | +0.95(%) | -1.18(%) |

### Dengue cases peak in late monsoon

Figure 2 depicts box and whisker plots, calculated for each month, of dengue cases based on eleven years, 2012 – 2022, of dengue data. The distribution of dengue cases varies across months. Number of dengue cases starts increasing after the arrival of monsoon season in Ahmedabad i.e. from the month of July. The maximum number of dengue cases are observed in the months of September and October. Dengue incident start declining from November onwards and attain a minimum in February - March. The 95^th^ percentile caps indicates that there are episodic high values of dengue in the months of September - November and that such episodes might be termed as dengue outbreak episodes depending on the normal baseline values of dengue cases in previous years. Dengue outbreak episodes can be defined as when there is a sudden increase in the number of dengue cases in a short span of time which is not normal for the given geographical location and time of the year.

**Figure 2:**
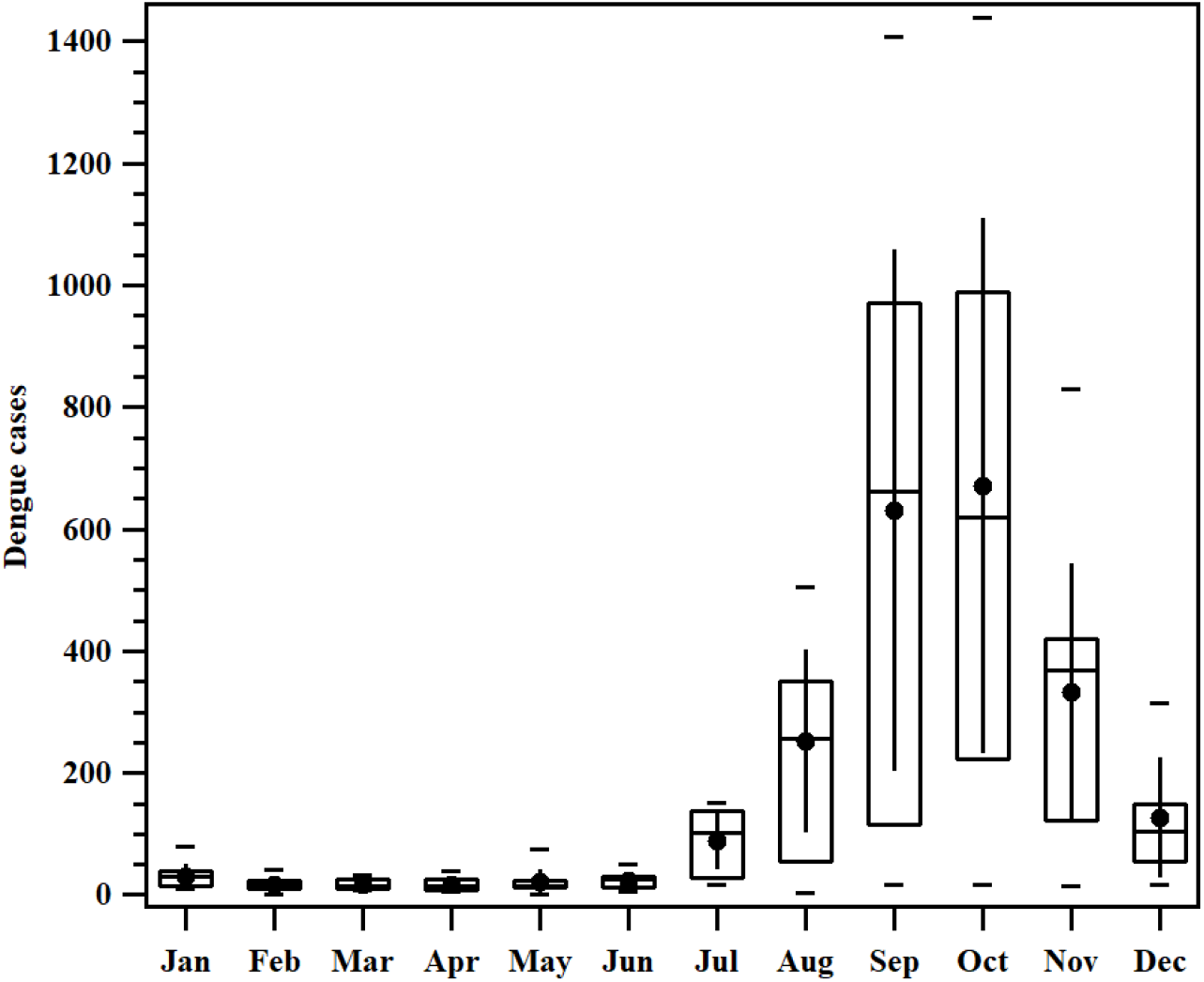
Monthly variation of dengue cases in the city of Ahmedabad based on data from 2012 – 2022. The symbols, lower whisker, lower edge of the box, central line of the box, upper edge of the box, upper whisker, solid circle, and vertical line represent 5^th^ percentile, 25^th^ percentile, median, 75^th^ percentile and 95^th^ percentile, mean, and standard deviation, respectively.

### Diurnal variations in temperature are minimum in certain months

Figure 3a shows the monthly variation of maximum and minimum temperature for the city of Ahmedabad based on eleven years, 2012 – 2022, of ERA5 reanalysis data. The city experiences the highest minimum temperature in the month of June while the monthly maximum temperature attains a peak in the month of May. Variability in both, the monthly maximum and minimum temperatures, reduces during the months of July to September. From July to September, both maximum and minimum temperatures show less variation. The difference between daytime highs and nighttime lows also shrinks from June to October, becoming smallest in August (Figure 3b). This means temperatures stay more consistent from day to day during these months. Overall, July to September bring a period of very stable temperature.

**Figure 3:**
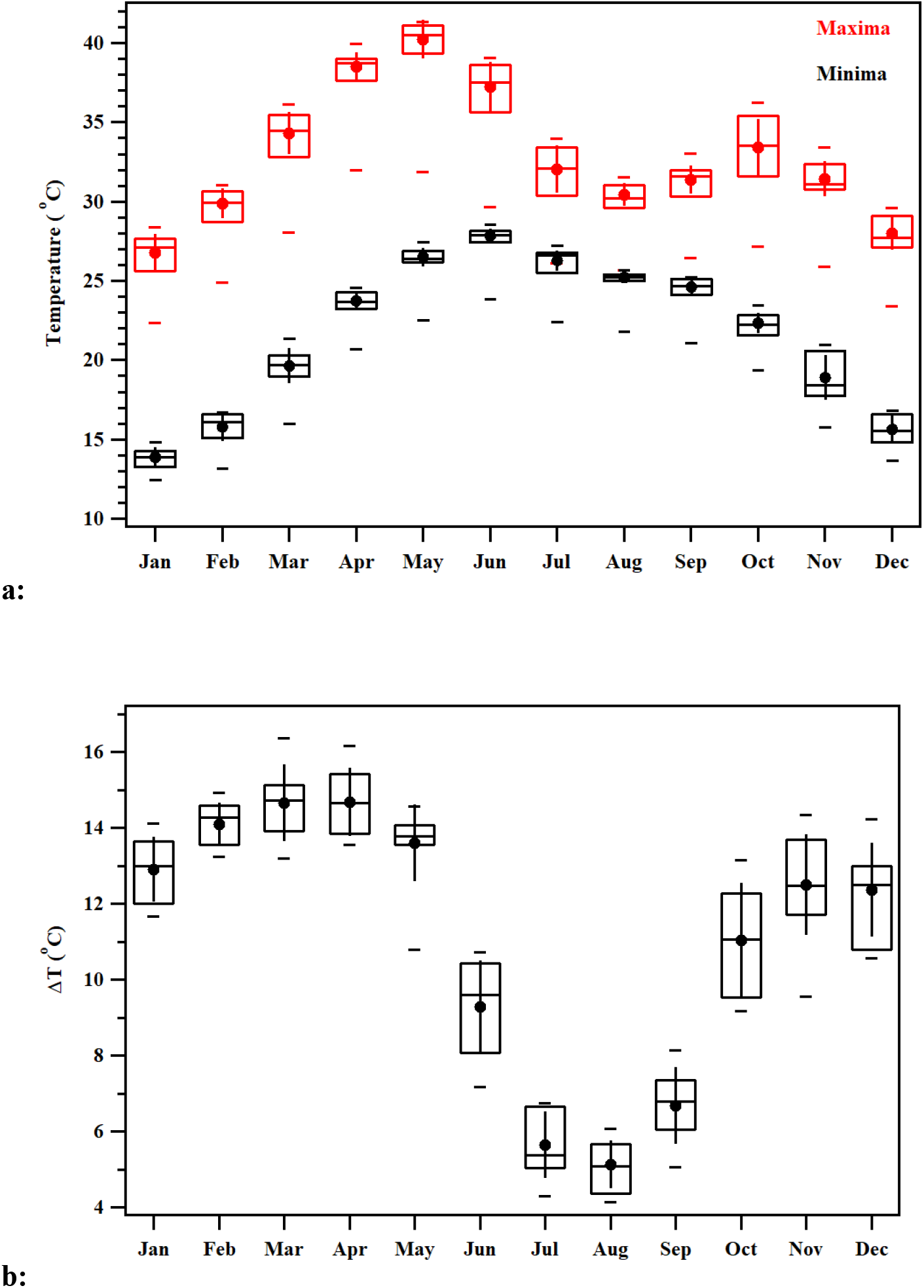
Monthly variation of a) maximum temperature (in red) and minimum temperature (in black), and b) the difference of monthly maximum and minimum temperature for the city of Ahmedabad based on eleven years, 2012 – 2022, of ERA5 reanalysis data. The symbols, lower whisker, lower edge of the box, central line of the box, upper edge of the box, upper whisker, solid circle, and vertical line are 5^th^ percentile, 25^th^ percentile, median, 75^th^ percentile and 95^th^ percentile, mean, and standard deviation, respectively.

### RH minimizes during post-monsoon months

Figure 4a shows the monthly variation of maximum and minimum RH for the city of Ahmedabad. Both, maximum and minimum RH, showed a bimodal feature where they peaked during the months of July - September and also peaked up during December-January. This is concomitant with the decrease of monthly variation of maximum temperature. The difference between maximum and minimum RH minimizes during the months of July - September thus, as seen in Figure 4b.

**Figure 4:**
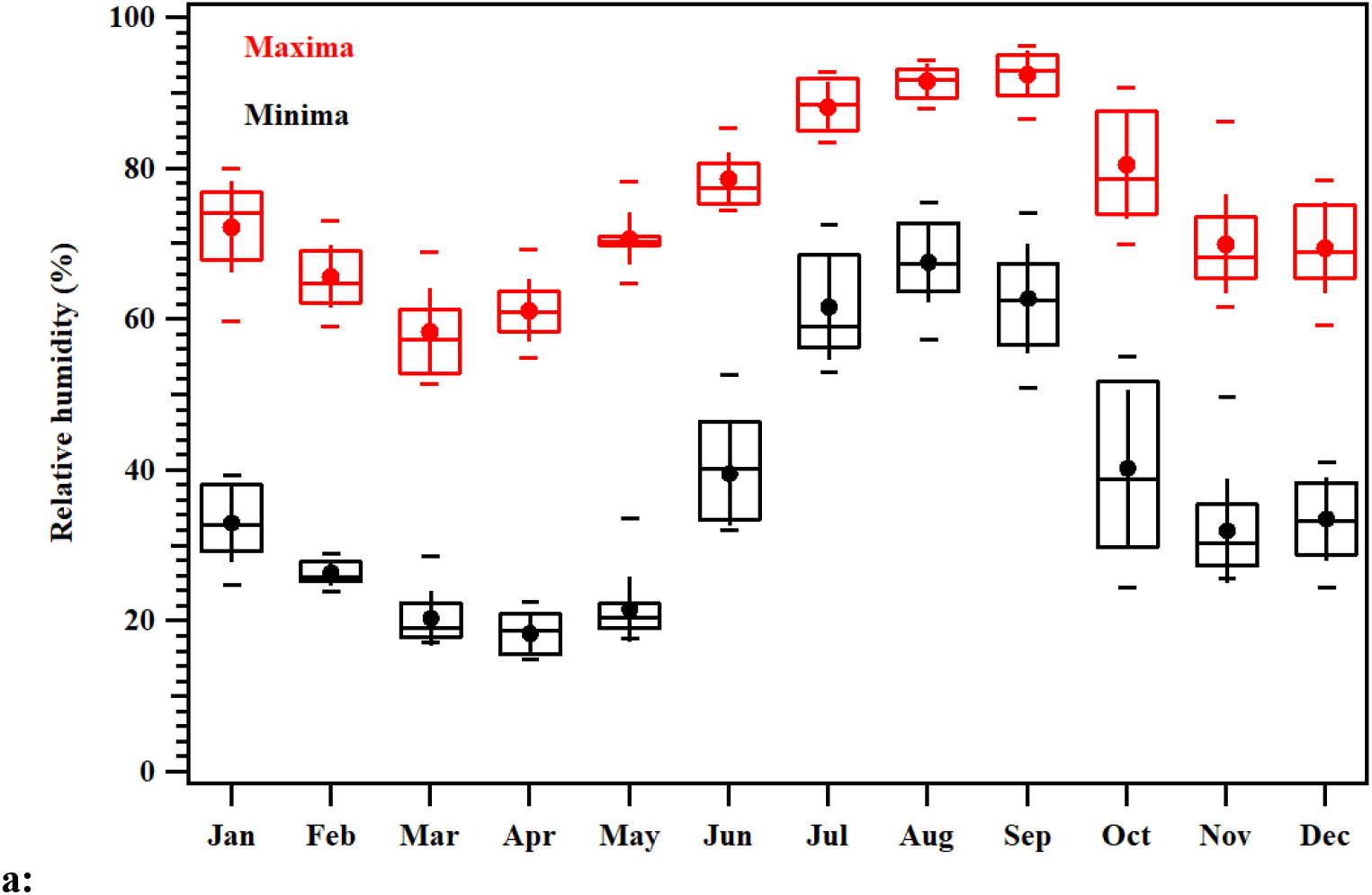
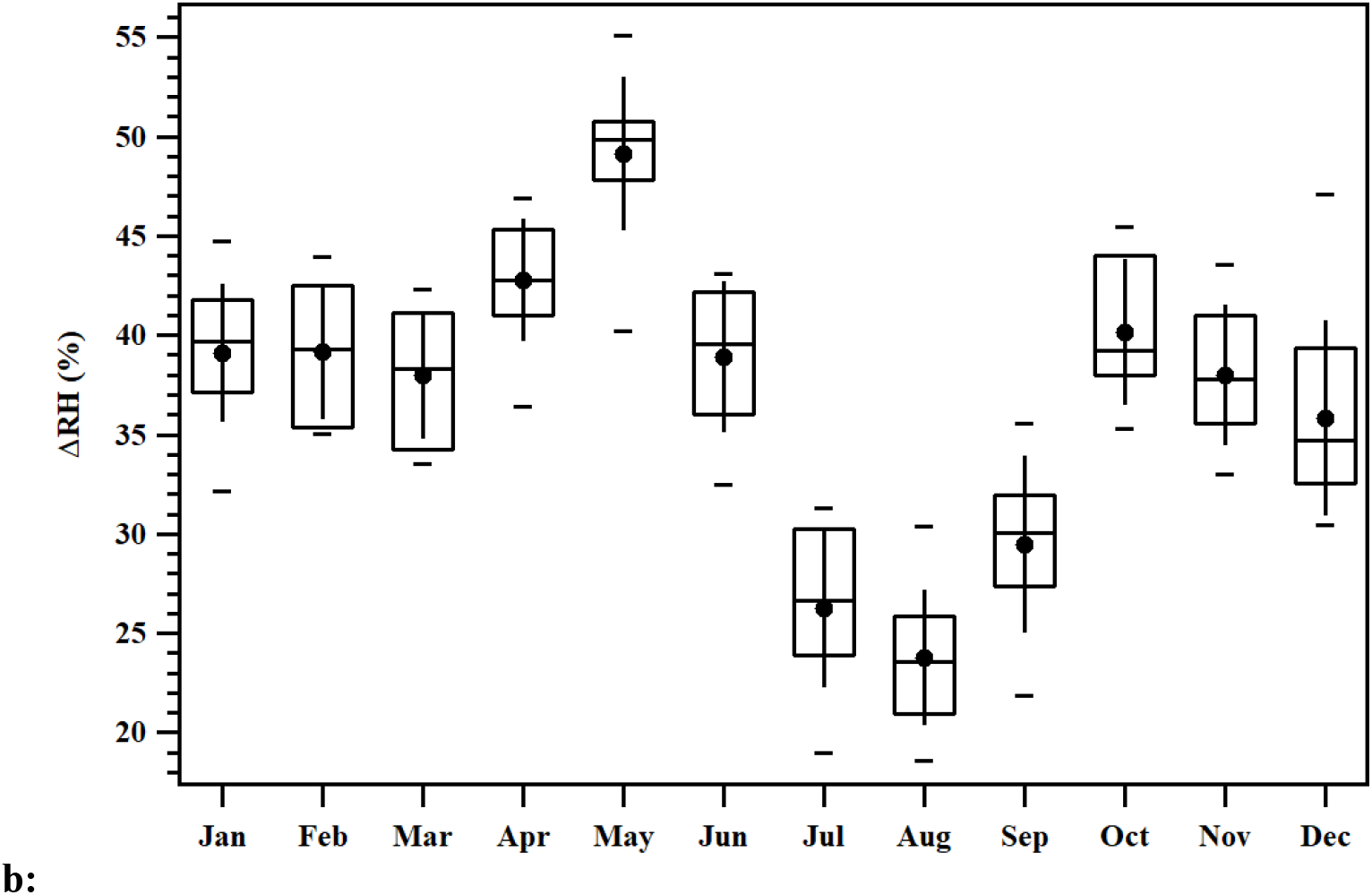
Monthly variation of a) maximum RH (in red) and minimum RH (in black), and b) the difference of maximum and minimum RH for the city of Ahmedabad based on eleven years, 2012 – 2022, of ERA5 reanalysis data. The symbols, lower whisker, lower edge of the box, central line of the box, upper edge of the box, upper whisker, solid circle, and vertical line are 5^th^ percentile, 25^th^ percentile, median, 75^th^ percentile and 95^th^ percentile, mean, and standard deviation, respectively.

### Stable temperature and humidity regime are crucial

RH and temperatures change diurnally and across days. We are interested in knowing specific interaction patterns of temperature and humidity changes where the number of dengue cases also go high or low in numbers. We assigned dengue cases of each month to a particular combination of temperature and RH bin having the month’s average temperature and RH range in that bin. A total of 132 data points of dengue (11 years) were assigned in this manner. The bin counts were additive in nature. Figure 5a shows occurrence of dengue cases as a function of binned average temperature and binned RH, all variables are based on monthly statistics. The highest number of dengue cases corresponds to 80-90% of RH and 25-30 °C temperatures. Figure 5b shows dengue cases plotted against the difference between maximum and minimum temperature (ΔT) and difference between maximum and minimum RH (ΔRH). While temperature triggers the dengue cases, RH drives the explosive phases and sustainability of the episode [63, 64] (Figure 5).

**Figure 5:**
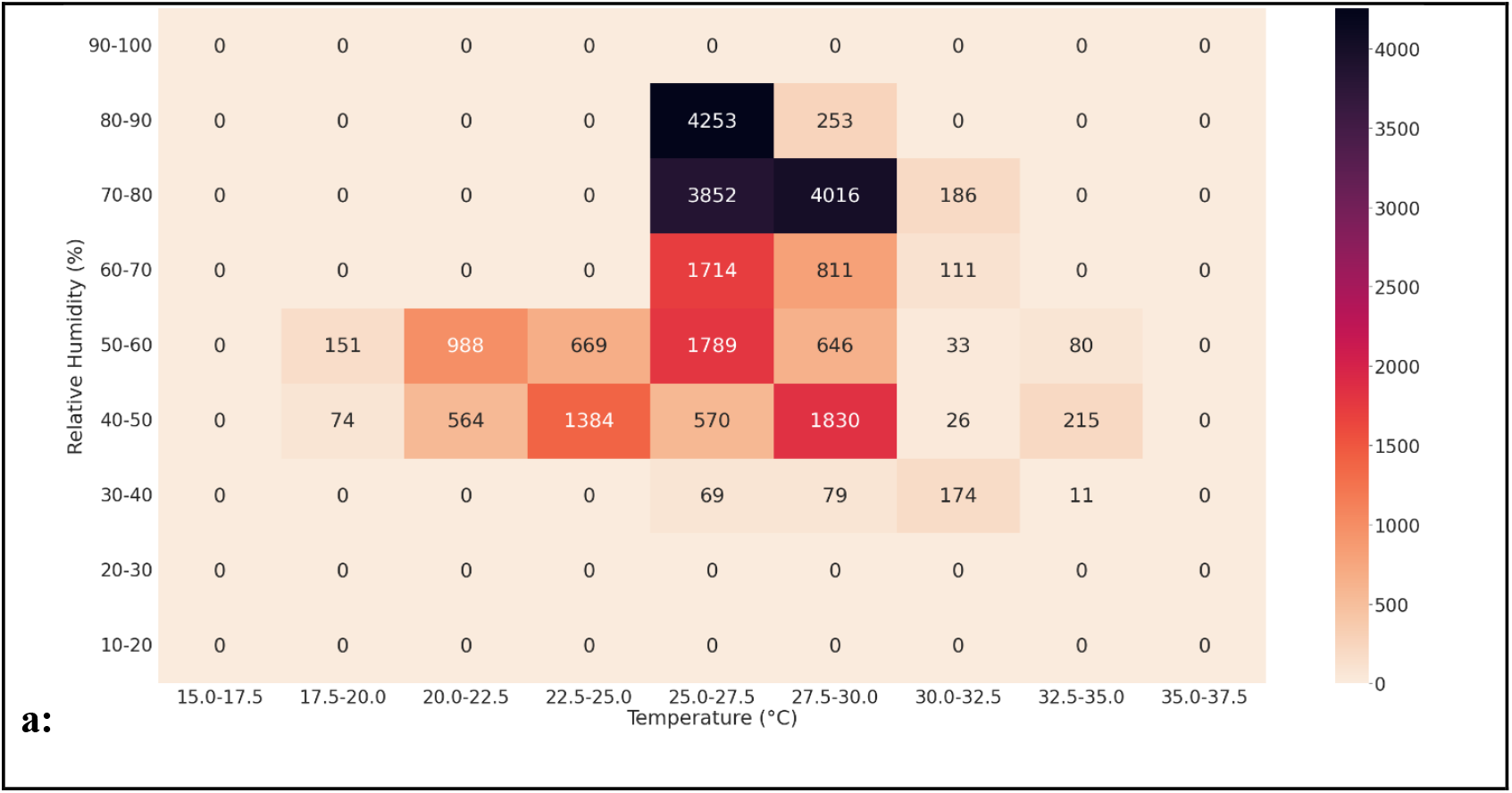
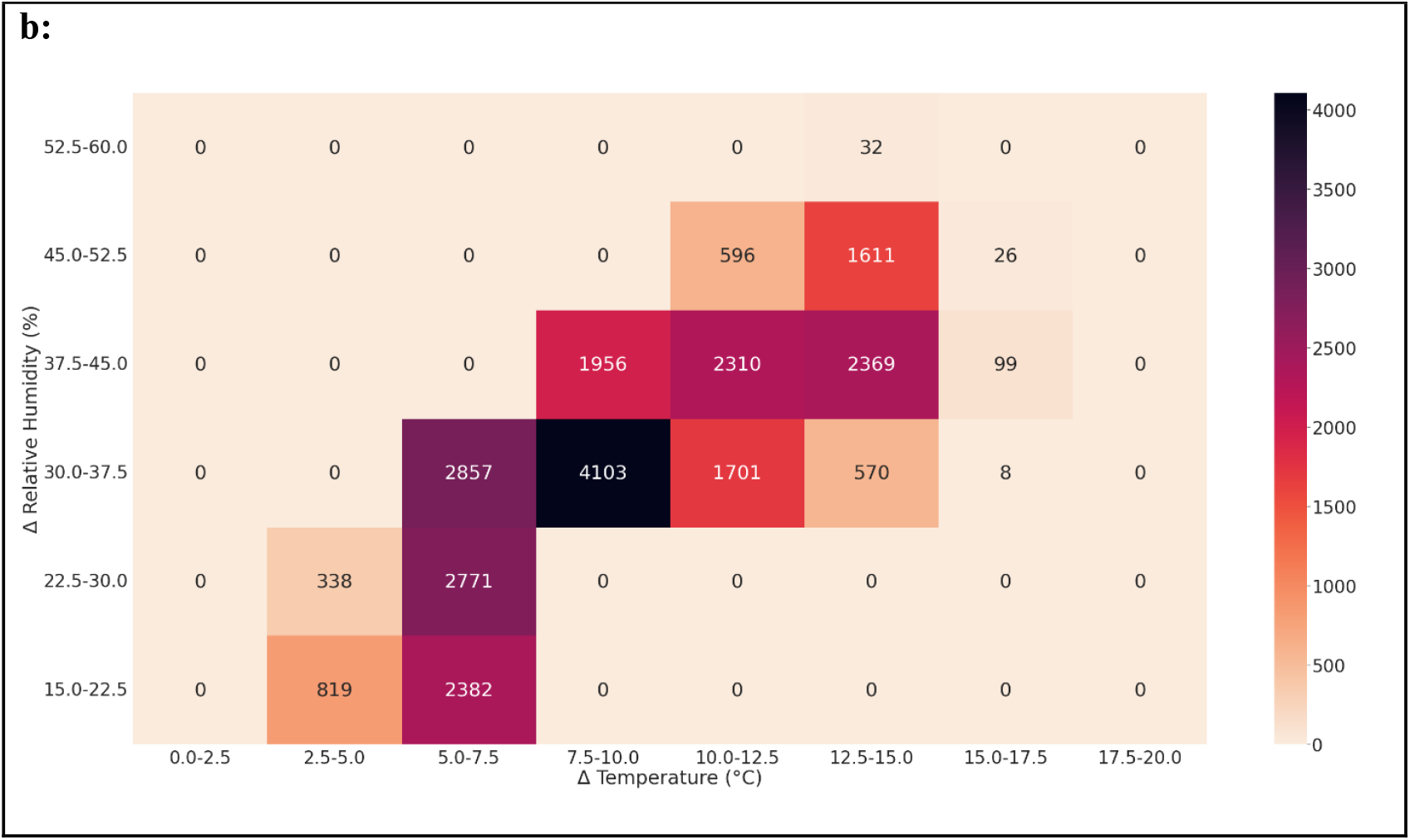
Occurrence of dengue cases as a function of a) binned average temperature and average RH, and b) binned differences in maximum temperature and minimum temperature (ΔT) and differences in maximum RH and minimum RH (ΔRH) within a month. The color bar represents the number of dengue cases. Dengue cases of each month were assigned to a bin having the month’s average temperature and RH range in that bin. A total of 132 data points (11 years) were distributed in this manner.

Based on the role of temperature and RH on the proliferation of dengue cases, and after narrowing down on the temperature and RH ranges responsible for high dengue cases for the city of Ahmedabad, we mapped the specific combinations of land surface temperature (LST) and RH when the frequency of occurrence of dengue cases mere maximum in the city. Based on the temperature and RH ranges when dengue cases were high as inferred from Figure 5, we found the frequency of occurrence those combinations, for the eleven year period concurrent with the dengue data, of land surface temperature and RH for each grid over the city of Ahmedabad. The grids with higher frequency of occurrence of the particular temperature and RH combinations are likely to be grids were dengue cases will see a spike, given other urban parameters are same. The spatial map of frequency of occurrence (count %) of favorable combinations of land surface temperature (LST) and relative humidity (RH) when dengue cases are maximum is shown in Figure 6. As can be seen from the spatial map, eastern parts of the city are more prone to dengue outbreaks when compared to south-western part. This granulated information on possible hotspots of dengue outbreaks will help city civic bodies to plan and take appropriate prevention and mitigation steps.

**Figure 6:**
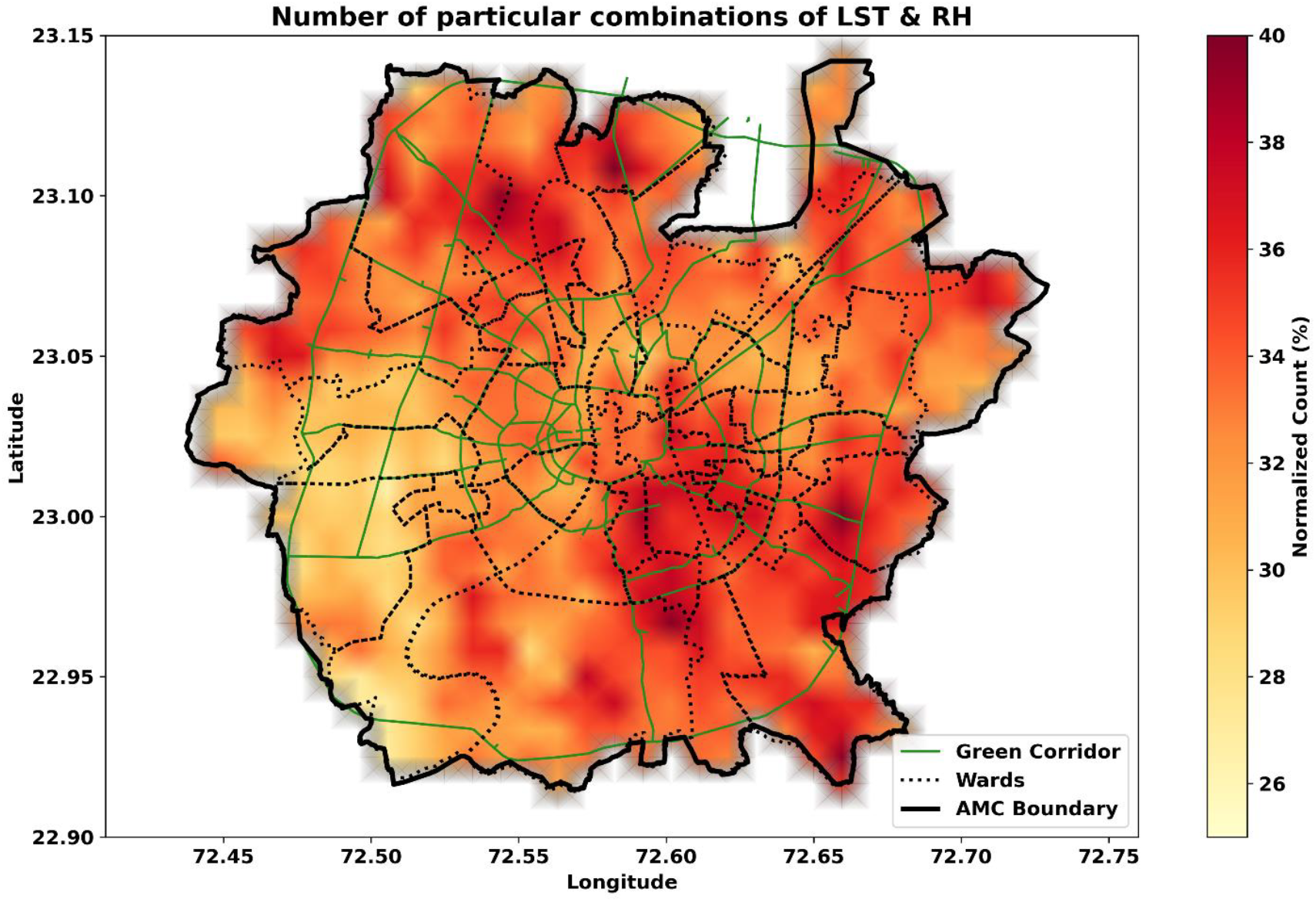
Spatial map of frequency of occurrence (count %) of favorable combinations of land surface temperature (LST) and relative humidity (RH) when dengue cases are maximum.

## Discussion

In Ahmedabad, the highest number of dengue cases, more than 600 case per month, occurred in September and October (Figure 2). This is concurrent with the monsoon period, and a stable temperature 28 °C - 32°C and RH 70 % -80 % regime. Mosquitoes have a narrow range of temperature and humidity windows that they can tolerate [65]. Delatte et al. [66] showed that egg hatching took the least time, approximately three days, at 20°C, while temperatures at 15 °C, 30 °C and 35 °C took around seven days [66]. The gonotrophic cycle was also calculated at different temperatures, and the shortest cycle, at an average of 3.5 days, was at 30°C [66]. This corresponds with the temperatures in Ahmedabad seen between July and October which are between 30 and 35 °C. The temperature, as well as the RH, are in the ideal range for the breeding of mosquitoes [67, 68]. At low humidity levels, they are sensitive to desiccation due to their high surface area-to-volume ratio. The longevity of *An. gambiae sensu stricto* (*s*.*s*.) was measured at 5°C intervals from 5°C to 40°C and with RH being 40%, 60%, 80%, and 100% [69]. With the RH being in the range of 60-100%, there was no significant difference in terms of survival. However, at 40% RH, the survival rate was seen to have decreased. Looking at Figure 3a, it can be seen that the maximum and minimum RH are still above 40% from June to October, which could play a role in the proliferation of *Aedes* species during the monsoon season.

### Non-fluctuating temperature-humidity interactions and dengue cases

Studies performed in other taxa (e.g. genus *Drosophila*) has evidenced higher egg laying in *D. melanogaster* females during the periods when certain combinations of temperature and humidity exist [52]. The temperature and humidity patterns observed in this study further emphasizes on these lines. Stable and minimal fluctuations in temperature and humidity sets ideal growth conditions for *A. aegypti* population expansion (indirect measures based on the dengue cases; Figure 5) as evidenced in our study (see Figures. 3B and 4B). Phenological shifts have also been observed in many insects and other ectotherms. During these phenological shifts, warming can increase development rates and the number of generations and benefiting to overall fitness [70]. The flip side is the invasion to novel sites. There are other factors which could play a role in the transmission and spread of DENV such as interspecies competition, parity, flight distances, urbanization as well as surveillance. Competition at the larval stage can also have an impact on the longevity of adult mosquitoes [71]. Moreover, a study conducted by Alto et al. [72, 73] showed that the interaction between adult mosquitoes and viral pathogens can be affected by larval competitions [72, 73]. At 27°C, 70% of female *Aedes aegypti* are parous and have a life expectancy of 35 days, based on a study conducted by Goindin et al. [74].

While our results are based on only two climatic variables it is based on a decade-long dengue data that was rigorously analysed in conjunction with these regional climatic variables. Dengue cases in India are clearly associated with monsoon patterns, and outcomes from the present analysis, specifically in terms of temperature and humidity bands, set a baseline to further develop predictive models. Our finding specifically emphasises the importance of delta values of temperature and humidity, which clearly associate with the breeding period of vectors, where a stable temperature and humidity ranges are crucial for the developmental growth and population expansion of vectors (i.e. Aedes aegypti). Some of the limitations of the study were due to dengue data resolution which was compiled at the municipal corporation repository on a monthly basis. This was one of the constraints in establishing a stronger association between dengue spike cases and climatic variables. This study entirely depended on the primary data sources and their resolution.

## Conclusions

Since dengue outbreak is a seasonal phenomenon in the tropics and subtropics, it is concentrated around the monsoon and post-monsoon seasons. It is primarily necessary to understand the dengue virus–environment interactions. This study, for the first time, investigates eleven years of dengue cases data from a densely populated metropolitan area of Ahmedabad and examines the trends and association with temperature and humidity changes. While the general behaviour of dengue virus is available, we have grounded this understanding in a city using the microclimate data and cases of dengue. Our findings are crucial and suggest that stable temperature and humidity patterns of certain combinations are crucial for dengue outbreaks in emerging urban landscapes in Indian cities. The highlights from the study are as follows:

- A stable temperature window of 25-27.5 °C and RH above 60% provide conducive conditions for dengue proliferation.
- While stable temperature ranges triggers the dengue cases, RH drives the explosive phases and sustainability of such episodes.
- Statistically significant increasing trends in temperatures and narrowing down of the day-night temperature ranges provide more stable temperature regimes in a warming world thereby likely to extend the dengue season beyond the usual monsoon season.

The study also shows spatial variation in the temperature-RH bands suitable for the outbreak of dengue, which the local government should pay attention to. Hence, the study can assist the local government in taking preventive educational and mitigating actions to control outbreak of dengue using temperature and RH variables, which are easily available data from the Indian Meteorological Department sources. This gives a tool in the hands of the local government to predict real time the potential periods and spots for dengue outbreak.

### Future directions

Temperature has an impact on the ability of mosquitoes to fly at certain distances. Rowley & Graham [75] show that the optimal temperature for flight is 27°C, and flight is possible between the range of 10 and 35°C [75]. In addition, RH, when above 30%, had a negligible effect on the flight. The increase in the human population correlates with the increase in urbanization. In a recent study by Sethi and Vinoj [76] it was found that the city of Ahmedabad has the highest overall increase in night-time LST due to urbanisation (Table 1). Further, since the night time temperatures are increasing faster than the daytime temperatures, urban centres like Ahmedabad are likely to witness more stable temperature regimes in a warming world. Such stable temperature regimes are likely to extend beyond the usual monsoon season thereby prolonging the dengue season. Therefore, natural land being converted for human use could change the disease risk due to the effect of interactions between people, vectors, vertebrate hosts, and pathogens [4, 77-82]. According to the Census of India, the population of Ahmedabad in 2011 was 6.36 million, and is projected to be 8.85 million in 2024. Therefore, the increase in population leads to an increase in urbanization, which, in turn, could contribute to the increase in mosquito-borne diseases, particularly dengue. In summary, the connections between urbanisation, population growth, and dengue proliferation should be explored.

## Declarations

### Abbreviations

Not applicable

### Ethics approval and consent to participate

Not applicable

### Consent for publication

Not applicable

### Availability of data and materials

All the meteorological data used in this study is open source, and can be accessed from the data portals of ECMWF and MODIS. Dengue data can be sourced from the Ahmedabad Municipal Corporation, Ahmedabad, Gujarat.

### Authors’ contributions

Conceptualization: SR & AV

Data collection and curation: CS, VP, LSC, SR, YD, AV, & KJ

Analysis and visualization: AV, KJ, & SR

Writing: SR, VP, AV, YD, DM, KJ, and BA

Resources: AV, SR, DM, BA, and KJ

## Acknowledgements

VP and YD were supported through PhD fellowship from Ahmedabad University. We thank Ahmedabad Municipal Corporation, Ahmedabad, Gujarat for their help in accessing dengue data for the last 11 years. We are thankful to the European Centre for Medium Range Weather Forecast (ECMWF) for the provision of ERA5-Land reanalysis products, some of which were used in this study. We acknowledge the usage of the Land Surface Temperature (LST) data, a satellite data product provided by the MODIS instrument onboard Terra satellite.

